# AdaGeneBudget: Cell-Adaptive Gene-Token Allocation for Efficient Single-Cell Foundation Models

**DOI:** 10.64898/2026.08.06.743174

**Authors:** Dohee Kim, Uiwon Hwang

## Abstract

Single-cell foundation models (scFMs) represent each cell using sequences of gene-associated tokens, making embedding extraction increasingly costly as the number of cells and expressed genes grows. Existing input policies typically rely on fixed input budgets, with retained genes determined by random subsampling, model-native ranking, or a dataset-level highly variable gene (HVG) panel. However, they do not jointly determine, for each cell, which genes to retain and how many tokens to allocate.

We introduce **AdaGeneBudget**, a training-free gene-token selection method that combines each gene’s expression with reference-derived inverse detection frequency and retains the shortest ranked prefix that captures a target fraction of the cell’s expression-specificity score mass. The resulting cell-specific budget is bounded by minimum and maximum lengths, requires no cell-type labels, and leaves the pretrained backbone unchanged.

We evaluated AdaGeneBudget in a frozen-backbone inference setting using pretrained scGPT and Geneformer models on Kang and PBMC reference-mapping tasks, with an additional scPRINT comparison against its official HVG policy and an expressed-only HVG control. Across four scGPT and Geneformer backbone–dataset pairs, AdaGeneBudget substantially reduced mean gene-token counts and peak GPU memory while increasing embedding-extraction throughput by up to 4.63×. Despite this compression, it preserved native-level aggregate annotation utility and numerically outperformed token-matched random and Top-Expression baselines. AdaGeneBudget also preserved fine-grained and low-support cell identities and retained lineage-marker programs and stimulation-associated pathway genes under compression. In scPRINT, both HVG controls achieved higher annotation macro-F1, whereas AdaGeneBudget more faithfully preserved the stimulation-induced embedding direction. These results establish biologically informed, cell-adaptive gene-token allocation as a practical complement to architectural and systems-level efficiency methods for applying existing scFMs to new datasets. They also suggest that cell-adaptive input allocation may be a useful design principle for future models operating under finite token budgets.

## 1. Introduction

The scale of single-cell RNA sequencing (scRNA-seq) datasets has increased rapidly, with public resources now aggregating tens of millions of cells across tissues and diseases.^1^ This growth has enabled single-cell foundation models (scFMs) to learn transferable gene and cell representations from large single-cell transcriptomic corpora. Geneformer and scGPT were pretrained on approximately 30 million and more than 33 million profiles, respectively, while scFoundation and scPRINT scale to 50 million or more cells.^2–5^ Once pretrained, these models provide reusable cell-level embeddings for downstream analyses such as clustering, cell-type annotation, and batch integration. As scFMs are applied to increasingly large atlas- and cohort-scale datasets, repeatedly extracting embeddings from long per-cell gene-token sequences can become a practical bottleneck under finite GPU memory and computation.

Despite differences in tokenization and expression encoding, many transformer-based scFMs represent each cell as a sequence of gene tokens. The number of vocabulary-matched expressed genes can vary substantially across cells and datasets and reach thousands,^2,3,5^ while pairwise computation in standard dense self-attention scales quadratically with sequence length.^6^ These computational demands become particularly important at atlas or cohort scale: more cells increase the total workload and longer per-cell sequences increase computation and memory per cell. Cell subsampling changes the set of cells retained for analysis, and uniform random subsampling may underrepresent rare populations or incompletely capture transcriptomic heterogeneity.^7^ We instead retain every cell while reducing the number of gene tokens used to represent each cell.

Outside single-cell modeling, related approaches include instance-wise feature selection, sparsity-inducing feature selection, and dynamic Transformer token pruning.^8–10^ These approaches typically rely on learned task-dependent selection mechanisms or pruning modules within the model. In contrast, AdaGeneBudget selects gene tokens before the native input pipeline of a frozen scFM, without modifying the backbone or requiring additional training.

Existing scFMs use model-specific but largely fixed input-construction rules. Geneformer rank-orders detected genes and truncates long inputs,^2^ whereas the official scGPT embedding pipeline randomly subsamples nonzero gene tokens when its maximum sequence length is exceeded.^3,11^ scPRINT was pretrained with randomly sampled gene contexts^5^ and also supports a dataset-level HVG policy that selects a fixed panel of highly variable genes.^12^ Random subsampling is simple but does not explicitly prioritize genes by cell-specific relevance. HVG selection provides a practical global policy because it ranks genes by their standardized cell-to-cell variability within the reference dataset and yields a shared ranking and panel across cells rather than adapting them to each cell.^13^

More specialized single-cell feature-selection and targeted-panel methods also construct compact gene sets: SCMER selects features to preserve cellular manifold structure, geneBasis selects a targeted gene panel to preserve transcriptomic structure, and PERSIST learns gene panels for transcriptome reconstruction or task-specific prediction.^14–16^ These methods typically produce gene sets shared across cells for their respective objectives, whereas AdaGeneBudget performs cell-specific gene prioritization and budget allocation to reduce gene-token processing cost while preserving downstream utility in an existing frozen scFM. Moreover, a common maximum length does not determine how many tokens are sufficient for each cell: some cells may be represented by a compact gene set, whereas others may require a broader one. This motivates treating input construction as two coupled, cell-specific decisions—which genes to retain and how many tokens to allocate—while bounding the resulting budgets to avoid extreme input lengths.

We introduce **AdaGeneBudget**, a training-free gene-token selection method for pre-trained scFMs. As illustrated in Fig. 1, AdaGeneBudget combines reference-derived inverse detection-frequency weights with each cell’s expression values and retains the shortest ranked prefix that reaches a target score mass within budget bounds. Cells with concentrated score mass therefore receive shorter inputs, whereas cells with more diffuse score mass receive longer inputs. The method requires no cell-type labels, estimates gene prevalence from the reference cohort only, and passes selected genes through each backbone’s native ordering and expression-encoding pipeline. We evaluate input policies with frozen backbones under the same reference-mapping protocol, isolating the effects of input construction from task-specific parameter adaptation, and show that AdaGeneBudget reduces embedding-extraction cost while preserving annotation utility and biologically relevant signals under gene-token compression.

**Fig. 1.**
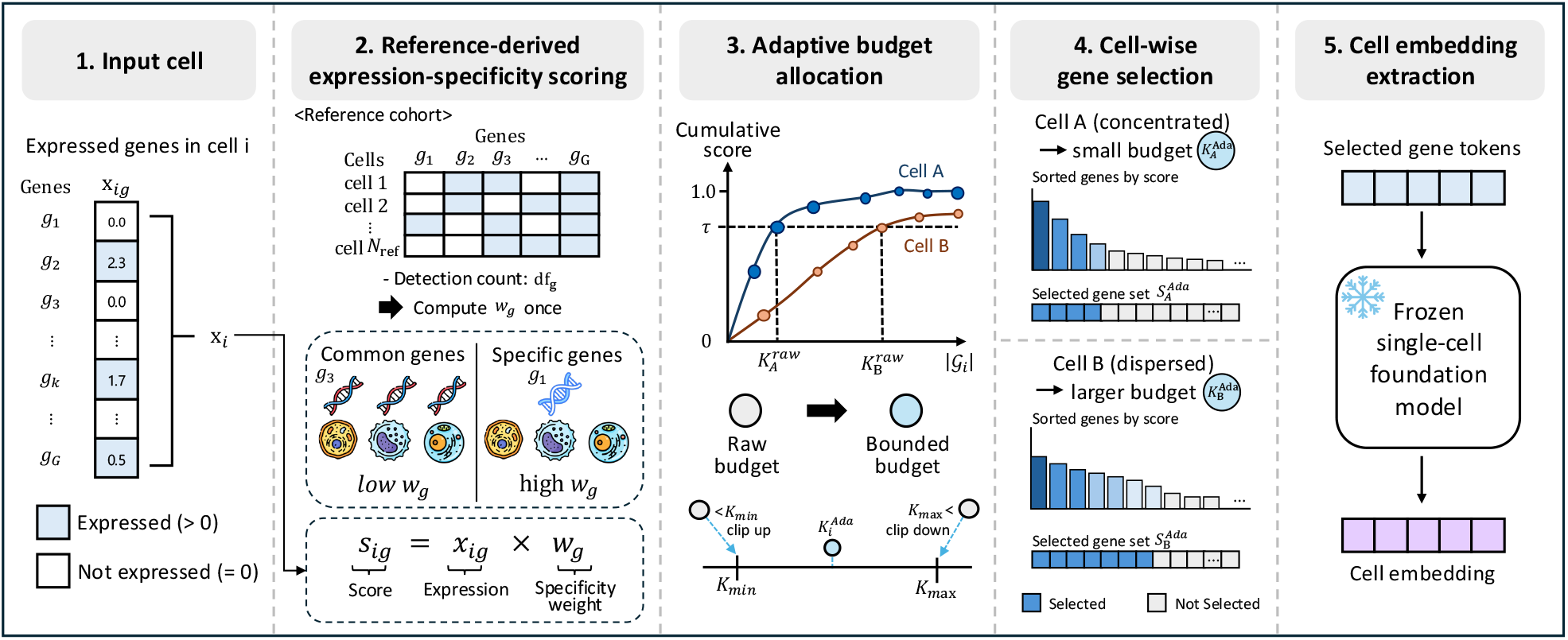
Overview of AdaGeneBudget. AdaGeneBudget computes reference-derived gene specificity, ranks expressed genes using expression-specificity scores, and assigns a cell-specific token budget according to cumulative score mass before constructing the native model input.

Our contributions are as follows. First, we formulate gene-token input construction in scFMs as two coupled cell-specific decisions: which genes to prioritize and how many tokens to allocate to each cell, without model retraining. Second, across scGPT and Geneformer, we demonstrate computational savings while preserving reference-mapping utility, using the fixed-budget Top-Expression baseline and the Fixed TF–IDF ablation to help distinguish the contribution of gene prioritization from that of adaptive budget allocation. Third, we show that its benefits extend beyond aggregate annotation, including preservation of fine-grained cell identities, lineage-marker programs, and stimulation-associated signals. Code and supplementary materials are available at https://github.com/D0heeKim/AdaGeneBudget.

## 2. Method

### 2.1. Problem formulation

For cell *i*, let 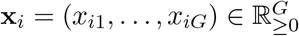 denote its expression vector, where *x*_*ig*_ is the expression value used to score gene *g*. Let *V* ⊆ {1, … , *G*} denote the set of genes recognized by the pretrained backbone. The eligible expressed-gene set and its size are

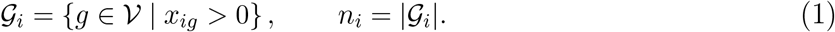

Under native input construction, each backbone constructs its model-specific input sequence from G_*i*_ according to its gene ordering, expression encoding, and input-length conventions. Our objective is instead to select a subset *S*_*i*_ ⊆ *G*_*i*_, with *K*_*i*_ = |S_*i*_|, to reduce gene-token processing cost while preserving downstream utility and biologically relevant information in the resulting frozen-backbone representation.

### 2.2. Reference-derived expression-specificity score

Expression magnitude captures within-cell abundance but not how frequently a gene is detected across the reference population. We therefore combine each gene’s expression value with an inverse reference-detection-frequency weight, following classical inverse-frequency weighting principles from information retrieval,^17,18^ yielding a TF–IDF-inspired expression-specificity score that reflects within-cell expression and inverse prevalence across the reference cohort.

Let the reference cohort contain *N*_ref_ cells. The reference detection count of gene *g*, analogous to document frequency, is

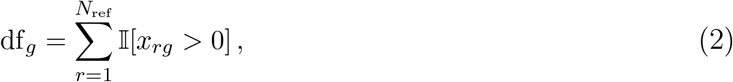

where I[·] denotes the indicator function. We then define the smoothed inverse-frequency weight for gene *g* and its expression-specificity score in cell *i* as

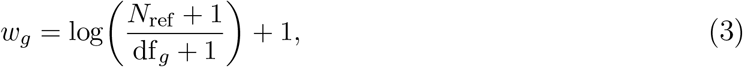

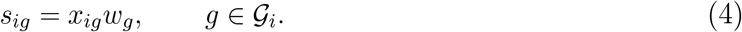

As df_*g*_ increases, *w*_*g*_ decreases toward 1; genes detected in many reference cells are therefore upweighted less, although they may still rank highly when *x*_*ig*_ is large.

We refer to *s*_*ig*_ as an *expression-specificity score*, rather than standard TF–IDF, because *x*_*ig*_ represents single-cell expression rather than textual term frequency. The weight vector is estimated once from the reference cohort and reused for both reference and held-out query cells. For each query cell, *s*_*ig*_ is computed from its own expression values using the fixed reference-derived weights; query-cohort prevalence and cell-type labels from either cohort are not used for ranking or selection.

### 2.3. Cell-adaptive score-mass allocation

For each cell *i*, let 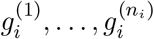 denote the eligible genes ordered by decreasing expression-specificity score:

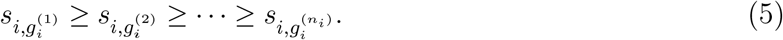

Ties are resolved deterministically according to each backbone’s fixed gene order. The normalized score mass captured by the top *k* genes is

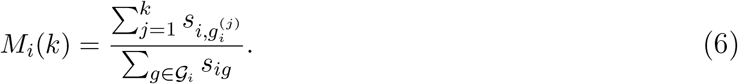

For a target score-mass threshold *τ* ∈ (0, 1], the unconstrained cell-specific budget is the length of the shortest prefix whose score mass reaches that threshold. Given 1 ≤ *K*_min_ ≤ *K*_max_, the unconstrained budget, the realized budget, and the selected gene set are defined as

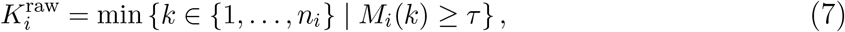

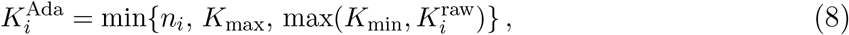

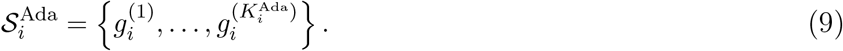

A cell with concentrated score mass reaches *τ* with a shorter prefix, whereas more diffuse score mass requires a longer prefix within the budget bounds. All main experiments use *τ* = 0.90, *K*_min_ = 128, and *K*_max_ = 600. Before applying the bounds, setting *τ* = 0.90 yields the shortest prefix that captures at least 90% of the cell’s total expression-specificity score mass; it does not retain a fixed proportion of the expressed genes. Sensitivity analyses are reported in Supplementary Sec. S1.

### 2.4. Backbone integration and computational implications

AdaGeneBudget operates before backbone-specific sequence construction and changes only the gene subset supplied to the backbone. Let *T*_*m*_ denote the sequence-construction function for backbone *m*, which maps the selected genes and their associated expression information to the model-specific input sequence, and let 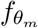 denote the corresponding embedding function. The resulting cell embedding is

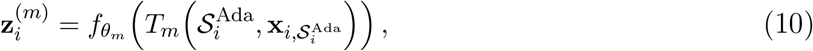

where 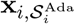 denotes the expression values of cell *i* restricted to the selected genes. After selection, each backbone’s native conventions for gene ordering, expression encoding, and special-token handling are retained; model-specific special tokens are not counted in 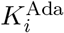. All backbone parameters remain fixed during embedding extraction. This separation allows AdaGeneBudget to be applied to other gene-token-based scFMs whose input pipelines accept an externally specified gene subset.

Given precomputed reference-derived weights, expression-specificity scoring requires *O*(*n*_*i*_) time, and the deterministic full ranking used in our implementation requires *O*(*n*_*i*_ log *n*_*i*_) time. Let 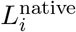 and 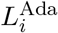 denote the realized unpadded input lengths, including model-specific special tokens, under native input construction and AdaGeneBudget, respectively. For the same backbone, reducing the input length reduces the pairwise token-interaction term in dense self-attention to approximately 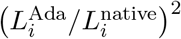 of its native value. This approximation applies only to the pairwise attention term; end-to-end runtime and memory also depend on selection overhead, token-wise computation, and batch padding. We therefore report mean realized gene-token count, end-to-end throughput, and peak GPU memory in Section 4.1, with selection overhead included in the throughput measurements.

## 3. Experimental setup

### Datasets and backbones

We evaluated AdaGeneBudget on Kang^19^ (19,583 reference cells from seven donors and 5,090 query cells from held-out donor 1015) and the PBMC RNA dataset from the Seurat v4 multimodal atlas^20^ (132,137 reference cells and 19,957 query cells from held-out donor P5), with both held-out splits fixed a priori based on dataset composition. The primary backbones were the scGPT whole-human checkpoint^3,11^ and Geneformer V1-10M.^2,21^ We additionally evaluated scPRINT on Kang because its dataset-level HVG option enables a same-backbone comparison between AdaGeneBudget and global HVG selection.^5,12^ We also audited evaluation-study overlap with documented pretraining corpora. PBMC over-laps with the source corpus used by scGPT, whereas no documented Kang source-study overlap was identified for scGPT or scPRINT. Kang and PBMC were absent from the documented Genecorpus-30M used by Geneformer V1. To assess transfer beyond the two primary immune-cell datasets, we additionally evaluated AdaGeneBudget on a triple-negative breast cancer (TNBC) dataset.^22^ Experimental details and results are provided in Supplementary Sec. S5.

### Input policies and evaluation

We compared AdaGeneBudget with Native Bucketed, Matched Random, and Top-Expression across both backbones, and additionally with Native Rank for Geneformer. Native Bucketed used each backbone’s native input policy with the same length-aware bucketing applied to all methods. Matched Random uniformly sampled eligible expressed genes without replacement using a fixed budget matched to AdaGeneBudget’s mean realized budget on the reference cohort. Top-Expression retained, for each cell, the highest-expressed eligible genes using the same reference-fitted fixed budget. For Geneformer, Native Rank retained genes in the backbone’s native rank-value order using the same fixed budget. Fixed budgets were matched to AdaGeneBudget’s mean realized budget on the reference cohort and then held fixed for query cells, without using query-set information for budget selection. For scPRINT, reference-only global HVG controls were computed using the Seurat v3 procedure;^13^ the complementary budget-matching schemes are described in Sec. 4.4.

Reference and query cells were embedded under each policy, and query cells were annotated by cosine 5-nearest-neighbor majority voting over the labeled reference embeddings; query labels were used only as ground truth for evaluation. Throughput included method-specific selection and bucketing but excluded shared data and model loading. Measurements were repeated five times, speedups were computed within each backbone–dataset pair, and all inference experiments used an NVIDIA RTX A6000 GPU.

### Biological-signal preservation metrics

PBMC lineage programs comprised 20 donor-consistent markers per lineage derived from the reference donors. Marker coverage was the fraction of a cell’s expressed lineage-program genes retained, and enrichment was coverage divided by the uniform-retention expectation *K*_*i*_*/n*_*i*_. Cells lacking expressed genes from their corresponding program were excluded, and cell-level values were averaged within each lineage.

For Kang, paired-donor pseudobulk differential expression was performed in CD14+ monocytes from the seven reference donors, and genes with *q* < 0.05 and | log_2_ FC| ≥ 1 were tested for one-sided Fisher over-representation in MSigDB Hallmark gene sets.^23^ After Benjamini-Hochberg correction,^24^ the 12 pathways with the smallest adjusted *P* values were selected for analysis. For each significant DE gene belonging to a selected pathway, mapped to the Geneformer input space and expressed in at least one held-out CD14+ monocyte, conditional retention was the fraction of expressing control and stimulated cells retaining that gene, and pathway retention was the unweighted mean across genes.

For scPRINT, Hallmark IFN-*α* response retention was the pooled fraction of expressed gene-cell pairs retained as positive input tokens in stimulated query cells.^23^ For cell type *c* and input policy *m*, perturbation-direction preservation was the cosine similarity between 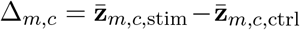 and the corresponding all-expressed reference direction from the same backbone, averaged equally across cell types with at least 20 query cells per condition.

## 4. Results

### 4.1. AdaGeneBudget reduces gene-token processing cost

Table 1 summarizes query-cell mean selected-gene counts, end-to-end embedding-pipeline throughput, and peak GPU memory, with the same length-aware bucketing applied to all policies and comparisons restricted within each backbone–dataset pair. Across the four backbone– dataset pairs, AdaGeneBudget reduced mean gene-token counts by 27.6–69.1%, yielding 1.19– 4.63× speedups and 42.5–70.6% lower peak memory than Native Bucketed.

**Table 1.** Computational efficiency. Speedup is relative to Native Bucketed within each backbone— dataset pair. Fixed-budget matching used reference cells, so query-set mean lengths may differ.

| Backbone | Dataset | Input policy | Mean genes ↓ | Throughput (cells/s) ↑ | Speedup | Peak memory (GB) ↓ |
|---|---|---|---|---|---|---|
| scGPT | Kang | Native Bucketed | 564.1 | 1129.9 ± 2.2 | 1.00× | 1.376 |
|  |  | Matched Random | 391.9 | 1438.7 ± 3.4 | 1.27× | 0.594 |
|  |  | Top-Expression | 391.9 | 1447.8 ± 38.5 | 1.28× | 0.594 |
|  |  | **AdaGeneBudget** | 405.8 | 1349.5 ± 2.1 | 1.19× | 0.791 |
|  | PBMC | Native Bucketed | 1184.7 | 443.9 ± 0.8 | 1.00× | 1.376 |
|  |  | Matched Random | 598.3 | 1031.2 ± 0.7 | 2.32× | 0.788 |
|  |  | Top-Expression | 598.3 | 990.2 ± 3.1 | 2.23× | 0.788 |
|  |  | **AdaGeneBudget** | 596.3 | 940.7 ± 5.1 | 2.12× | 0.791 |
| Geneformer | Kang | Native Bucketed | 558.3 | 810.7 ± 7.4 | 1.00× | 13.527 |
|  |  | Native Rank | 390.3 | 1243.7 ± 0.5 | 1.53× | 2.654 |
|  |  | Matched Random | 390.3 | 1199.0 ± 0.4 | 1.48× | 2.654 |
|  |  | Top-Expression | 390.3 | 1192.2 ± 8.8 | 1.47× | 2.654 |
|  |  | **AdaGeneBudget** | 404.2 | 1101.6 ± 0.4 | 1.36× | 3.981 |
|  | PBMC | Native Bucketed | 1927.5 | 146.7 ± 2.7 | 1.00× | 13.527 |
|  |  | Native Rank | 598.2 | 721.6 ± 2.1 | 4.92× | 3.971 |
|  |  | Matched Random | 598.2 | 744.1 ± 1.0 | 5.07× | 3.971 |
|  |  | Top-Expression | 598.2 | 716.1 ± 4.9 | 4.88× | 3.971 |
|  |  | **AdaGeneBudget** | 596.1 | 678.6 ± 0.3 | 4.63× | 3.977 |

Matched Random was 6.6–9.7% faster because it avoided expression-specificity scoring, full ranking, and adaptive cutoff determination. On Geneformer–Kang, AdaGeneBudget selection accounted for 14.8% of the end-to-end embedding-pipeline time; within selection, the TF–IDF and native-order sorts dominated the profiled cost (68.3%) compared with 12.2% to scoring and 11.2% to cutoff determination. Nevertheless, AdaGeneBudget retained 91.2–93.8% of Matched Random throughput. Its higher peak memory than fixed-budget policies with similar mean lengths reflects the longest padded batches, as variable allocation allows cells to reach *K*_max_.

### 4.2. AdaGeneBudget preserves reference-mapping annotation utility under compression

Under the reference-mapping protocol in Section 3, AdaGeneBudget numerically outperformed Matched Random and Top-Expression on all three annotation metrics and all four backbone– dataset pairs (Table 2), with macro-F1 gains of 2.24–18.04 percentage points over Matched Random. Despite substantially compressed inputs, it also numerically exceeded Native Bucketed on all three metrics across all four backbone–dataset pairs and Geneformer Native Rank in both datasets, indicating that which genes are retained matters, not only how many.

**Table 2.**
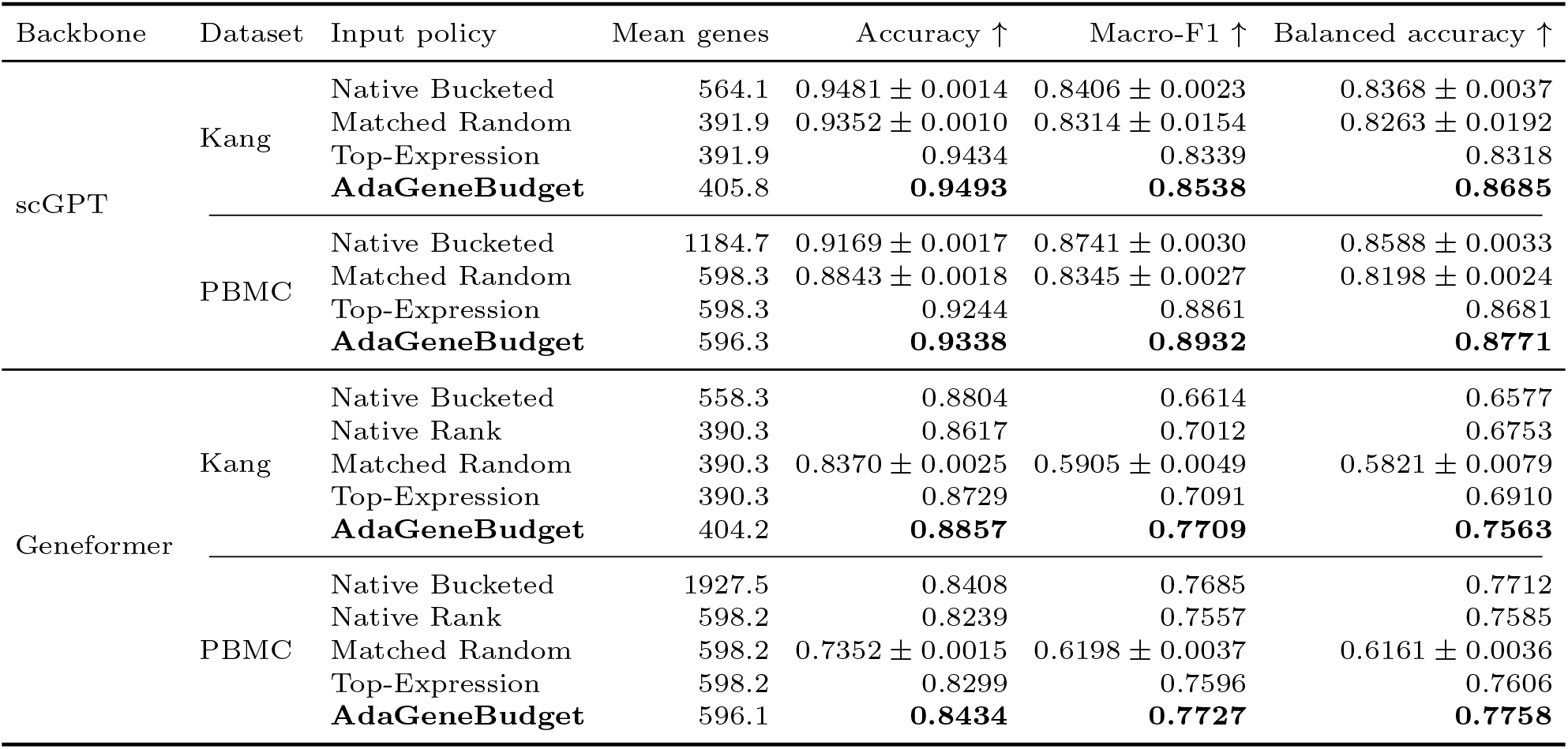
Reference-to-query cell-type annotation performance. Stochastic policies are reported as mean ± standard deviation across selection seeds, whereas deterministic policies are reported once. Bold indicates the best numerical result within each backbone–dataset pair.

| Backbone | Dataset | Input policy | Mean genes | Accuracy ↑ | Macro-F1 ↑ | Balanced accuracy ↑ |
|---|---|---|---|---|---|---|
| scGPT | Kang | Native Bucketed | 564.1 | 0.9481 ± 0.0014 | 0.8406 ± 0.0023 | 0.8368 ± 0.0037 |
|  |  | Matched Random | 391.9 | 0.9352 ± 0.0010 | 0.8314 ± 0.0154 | 0.8263 ± 0.0192 |
|  |  | Top-Expression | 391.9 | 0.9434 | 0.8339 | 0.8318 |
|  |  | **AdaGeneBudget** | 405.8 | **0.9493** | **0.8538** | **0.8685** |
|  | PBMC | Native Bucketed | 1184.7 | 0.9169 ± 0.0017 | 0.8741 ± 0.0030 | 0.8588 ± 0.0033 |
|  |  | Matched Random | 598.3 | 0.8843 ± 0.0018 | 0.8345 ± 0.0027 | 0.8198 ± 0.0024 |
|  |  | Top-Expression | 598.3 | 0.9244 | 0.8861 | 0.8681 |
|  |  | **AdaGeneBudget** | 596.3 | **0.9338** | **0.8932** | **0.8771** |
| Geneformer | Kang | Native Bucketed | 558.3 | 0.8804 | 0.6614 | 0.6577 |
|  |  | Native Rank | 390.3 | 0.8617 | 0.7012 | 0.6753 |
|  |  | Matched Random | 390.3 | 0.8370 ± 0.0025 | 0.5905 ± 0.0049 | 0.5821 ± 0.0079 |
|  |  | Top-Expression | 390.3 | 0.8729 | 0.7091 | 0.6910 |
|  |  | **AdaGeneBudget** | 404.2 | **0.8857** | **0.7709** | **0.7563** |
|  | PBMC | Native Bucketed | 1927.5 | 0.8408 | 0.7685 | 0.7712 |
|  |  | Native Rank | 598.2 | 0.8239 | 0.7557 | 0.7585 |
|  |  | Matched Random | 598.2 | 0.7352 ± 0.0015 | 0.6198 ± 0.0037 | 0.6161 ± 0.0036 |
|  |  | Top-Expression | 598.2 | 0.8299 | 0.7596 | 0.7606 |
|  |  | **AdaGeneBudget** | 596.1 | **0.8434** | **0.7727** | **0.7758** |

A Fixed TF–IDF ablation using the same score and ranking as AdaGeneBudget but a shared fixed budget supported the contribution of adaptive allocation: on Geneformer–Kang, AdaGeneBudget increased macro-F1 from 0.7130 to 0.7709 under comparable realized budgets. The default configuration remained close to the best tested across all four backbone– dataset pairs (Supplementary Sec. S1). Supplementary Sec. S2–S4 show that larger macro-F1 gains were supported by bootstrap intervals, AdaGeneBudget better preserved local, within-cell-type, and expression-referenced structure than Matched Random and tested reference variations had limited effects on Geneformer–Kang.

Supplementary donor-disjoint validation on a 427,823-cell TNBC cohort further extended the efficiency–utility tradeoff beyond the two primary immune-cell datasets. AdaGeneBudget achieved 1.80× and 3.46× speedups over Native Bucketed with scGPT and Geneformer, respectively, while attaining the highest numerical macro-F1 among the evaluated policies for both backbones (0.5411 and 0.3392; Supplementary Sec. S5).

### 4.3. AdaGeneBudget preserves fine-grained and low-support cell identities

Aggregate annotation metrics can mask failures in fine-grained or low-support populations. We therefore examined Geneformer–Kang performance by L1 support and scGPT–PBMC performance using finer L2 labels (Fig. 2).

**Fig. 2.**
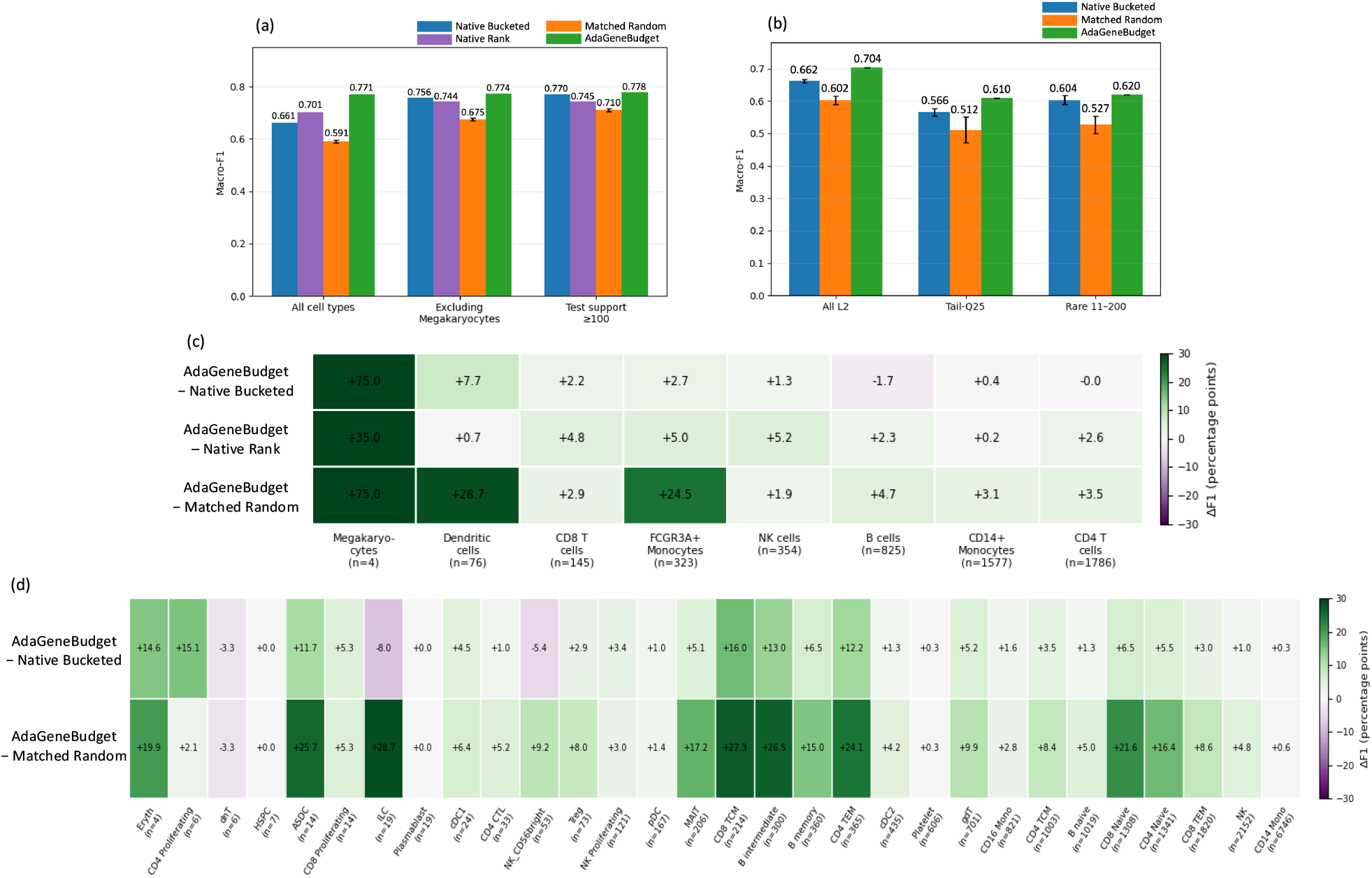
Fine-grained and low-support annotation under gene-token reduction. (a) Geneformer–Kang L1 macro-F1 across all cell types and support-restricted subsets. (b) scGPT– PBMC L2 macro-F1 across all subtypes and low-support subsets. (c,d) Per-population F1 differences, in percentage points, between AdaGeneBudget and the indicated comparators for Geneformer–Kang (c) and scGPT–PBMC (d), ordered by support. Error bars show standard deviation across five seeds for stochastic policies; deterministic policies are single runs.

On Kang, AdaGeneBudget achieved the highest macro-F1 across all cell types (0.771), after excluding Megakaryocytes (0.774) and among classes with at least 100 query cells (0.778) (Fig. 2a,c). Thus, the aggregate gain was not driven by an extremely low-support population.

For Megakaryocytes (*n* = 4), AdaGeneBudget correctly annotated three cells (F1=0.750), compared with F1=0 for Native Bucketed and Matched Random and 0.400 for Native Rank; given the very small support, we treat this result as an illustrative example rather than independent evidence of rare-cell robustness.

On PBMC, AdaGeneBudget likewise achieved the highest macro-F1 across all 30 L2 sub-types (0.704), in Tail-Q25 (0.610), and among subtypes with 11–200 cells (0.620) (Fig. 2b). Among subtypes with more than 10 query cells, it exceeded Native Bucketed for 23 and Matched Random for 25, including all 11 T-cell-related subtypes (Fig. 2d). The only sub-types above 10-cell support for which AdaGeneBudget was lower than Native Bucketed were ILC and NK CD56^bright^, by 8.0 and 5.4 points, respectively, while remaining above Matched Random by 28.7 and 9.2 points. These exceptions may reflect the difficulty of distinguishing related lymphoid populations, since human ILC states are transcriptionally heterogeneous and some ILC1 and NK states show transcriptional similarity.^25,26^ Moreover, the original PBMC subtype labels were derived from joint RNA and surface-protein analysis,^20^ which may make exact recovery from RNA-only embeddings challenging. Among the four subtypes below the 11-cell support, only dnT was lower than both comparators (3.3 points); with only six cells, both Native Bucketed and Matched Random achieved mean F1 of 0.033, with a nonzero result in just one of five seeds for each policy. We therefore do not interpret this small difference as a stable comparator advantage. Overall, AdaGeneBudget preserved fine-grained and low-support identities across support-restricted analyses, with isolated exceptions confined to extremely small or closely related lymphoid populations.

### 4.4. AdaGeneBudget preserves lineage and stimulation-associated signals under compression

Beyond annotation utility, we evaluated whether compression retained stimulation-associated pathways, lineage-marker programs, and stimulation-induced embedding shifts (Fig. 3).

**Fig. 3.**
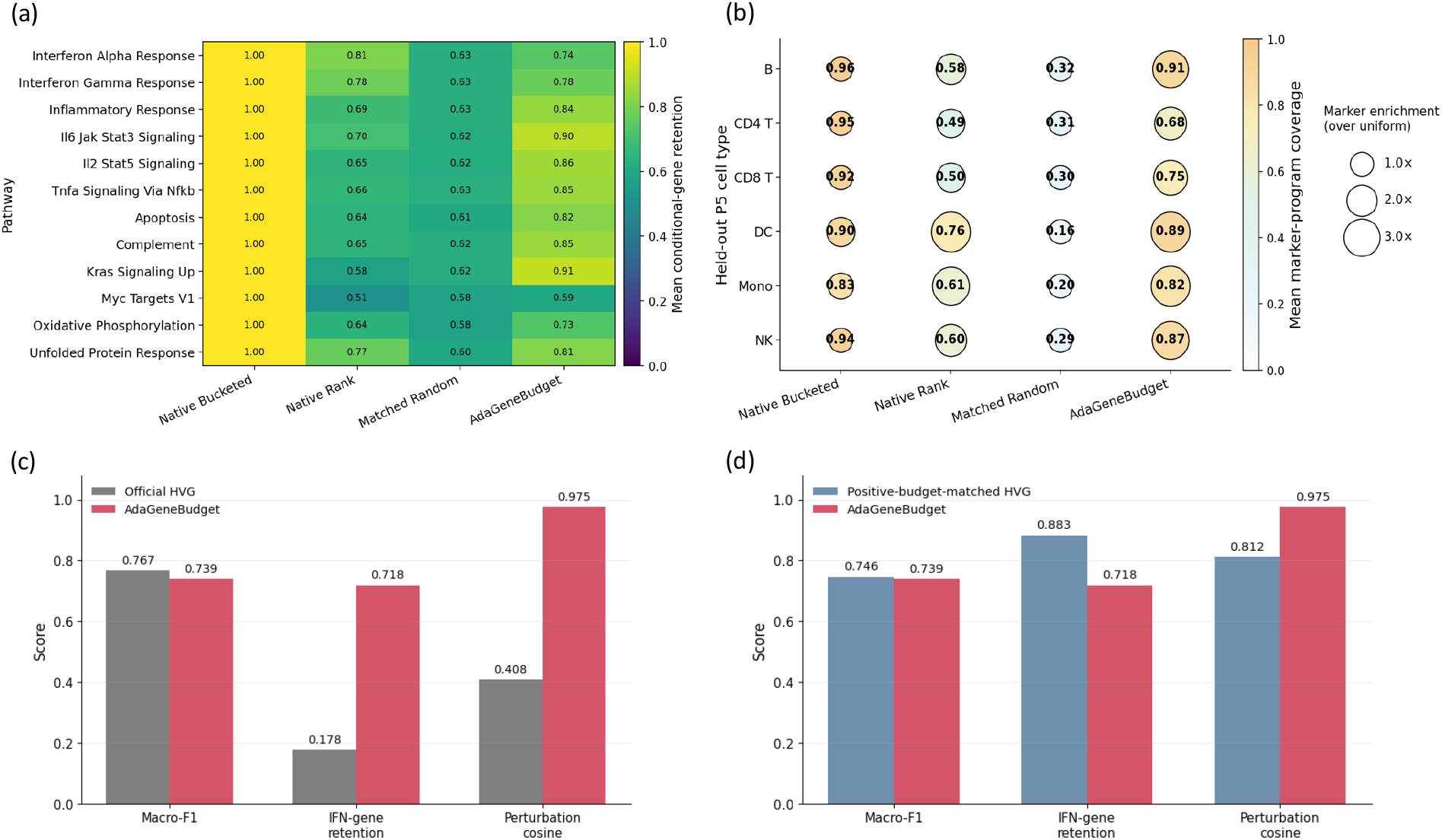
Biological-signal preservation under compression. (a) Geneformer–Kang mean conditional retention of stimulation-associated pathway genes. (b) Geneformer–PBMC lineage-marker coverage in held-out donor P5 (color and values) and enrichment over uniform retention at each policy’s realized budget (bubble area). (c) scPRINT’s reference-only Seurat-v3 fixed-panel HVG policy versus AdaGeneBudget under a matched total-token budget. (d) Expressed-only HVG versus Ada-GeneBudget under a matched positive-gene budget. Panels (c,d) report macro-F1, IFN-*α*-response retention, and cosine similarity to the all-expressed stimulation direction.

In Geneformer–Kang, AdaGeneBudget achieved mean conditional retention of approximately 0.81, compared with 0.67 for Native Rank and 0.61 for Matched Random (Fig. 3a). It exceeded Native Rank in 10 of 12 pathways and Matched Random in all 12, with high retention for IL6–JAK–STAT3, TNF*α*–NF*κ*B, and KRAS signaling.

In Geneformer–PBMC, AdaGeneBudget retained approximately 0.82 mean lineage-marker coverage versus 0.92 for Native Bucketed, 0.59 for Native Rank, and 0.26 for Matched Random (Fig. 3b). This recovered about 89% of native coverage while reducing mean input from 1,927.5 to 596.1 genes and exceeded comparators across all six lineages. Bootstrap uncertainty estimates for both biological-preservation analyses are provided in Supplementary Sec. S2.

With scPRINT, the official fixed-panel HVG control was matched to AdaGeneBudget’s mean total gene-token count on the reference cohort, including zero-valued panel entries. HVG achieved higher macro-F1 (0.767 versus 0.739), whereas AdaGeneBudget retained more Hallmark IFN-*α*-response genes (0.718 versus 0.178) and more faithfully preserved the stimulation-induced embedding direction (0.975 versus 0.408; Fig. 3c).

To separate the effect of zero-expression overhead, we also evaluated an expressed-only HVG control by expanding the reference-fitted global candidate panel to 11,929 genes to match AdaGeneBudget’s mean positive-input length (Fig. 3d). HVG achieved slightly higher macro-F1 (0.746 versus 0.739) and IFN-response retention (0.883 versus 0.718), whereas Ada-GeneBudget better preserved the stimulation direction (0.975 versus 0.812), with higher similarity in all seven eligible cell types. Thus, the direction-preservation advantage persisted even under this broader expressed-only HVG control.

## 5. Discussion

We introduced AdaGeneBudget as a training-free method that treats gene-token input construction in scFMs as a cell-specific allocation problem. Across scGPT and Geneformer, it reduced embedding-extraction cost while preserving native-level annotation utility; its advantage over token-matched random selection and Geneformer Native Rank indicates that representation quality depends on which genes are retained as well as how many. Our empirical conclusions concern frozen-backbone embedding extraction and reference mapping; we did not evaluate pretraining, fine-tuning, or task-specific model adaptation. Held-out donors were unseen only with respect to the reference-mapping split, not necessarily model pretraining: corpus audits confirmed PBMC overlap for scGPT and at the scPRINT source-corpus level, whereas no source-study match for Kang or PBMC was identified in the documented Geneformer V1 corpus. We therefore do not characterize all evaluated settings as pre-training-unseen.

The biological analyses further demonstrate why evaluation should extend beyond aggregate annotation metrics. Support-stratified analyses showed that gains over random compression persisted after excluding the extremely low-support Megakaryocyte population and among more highly supported cell types. AdaGeneBudget also retained lineage-marker programs and stimulation-associated pathway genes at rates far above random selection. In scPRINT, both HVG controls achieved higher annotation macro-F1, whereas AdaGeneBudget more faithfully preserved the stimulation-induced embedding direction under both fixed-panel and expressed-only comparisons. These findings suggest that biological-state fidelity should be considered alongside cell-type annotation when evaluating compressed scFM inputs.

AdaGeneBudget complements rather than replaces architectural and systems-level efficiency improvements. Because it operates before the native input pipeline, requires no backbone updates or cell-type labels, and computes reference-derived statistics once, it can be applied to frozen models without retraining. The same principle may also inform future scFM design: when candidate genes exceed an explicit or practical context budget, identical maximum lengths may not be the most effective use of computation.

Several limitations remain. First, the main evaluation focused on two immune-cell datasets and three pretrained backbones. Supplementary donor-disjoint TNBC validation broadened the tissue and disease settings with a 427,823-cell cohort, but broader validation across tissues, sequencing settings, and pretrained checkpoints is still needed. Evaluating whether similar benefits extend to pretraining or task-specific fine-tuning also remains future work. Second, the expression-specificity weights depend on the reference cohort. Although performance was stable under the tested changes in reference size, donor composition, and under cross-cohort weight estimation, robustness to broader biological or technical reference–query shifts remains to be established. Because inverse-frequency weighting favors low-frequency genes, rare technical artifacts may also receive increased priority. In scGPT–PBMC, low-frequency genes were enriched among selected inputs but remained a small absolute fraction; detailed results are provided in Supplementary Sec. S6. Third, the current scoring rule evaluates genes individually and does not explicitly account for redundancy among selected genes. Under a limited token budget, highly correlated or co-expressed genes may therefore occupy multiple input positions while providing overlapping information, potentially displacing genes with complementary signals. Finally, adaptive scoring and full ranking introduce additional preprocessing overhead relative to simpler fixed-budget policies. Sorting accounted for most of this cost, suggesting that partial-ranking or approximate top-*k* implementations could reduce it. Because runtime and memory were evaluated on a single GPU platform, the observed computational gains should not be assumed to transfer unchanged across hardware environments.

Overall, AdaGeneBudget shows that cell-adaptive, biologically informed token allocation can provide a favorable balance between computational efficiency and representation fidelity. This establishes input gene selection as a practical and complementary direction for scaling gene-token-based scFMs.

## 6. Acknowledgements

This work was supported by the National Research Foundation of Korea (NRF) grant funded by the Korea government (MSIT) (RS-2025-00561169), Global – Learning & Academic research institution for Master’s · PhD students, and Postdocs (G-LAMP) Program of the National Research Foundation of Korea (NRF) grant funded by the Ministry of Education (No. RS-2025-25442252), Institute of Information & communications Technology Planning & Evaluation (IITP) under the Leading Generative AI Human Resources Development (IITP-2027-RS-2026-25546026) grant funded by the Korea government (MSIT), and the NAVER Digital Bio Innovation Research Fund, funded by NAVER Corporation (Grant No. 3720260100).

