## Supplementary Material for "AdaGeneBudget: Cell-Adaptive Gene-Token Allocation for Efficient Single-Cell Foundation Models"

#### *S1. Ablation and Hyperparameter Analysis*

**Adaptive allocation and cell-specific budgets.** To isolate the contribution of adaptive budget allocation while holding the gene-ranking rule fixed, we compared AdaGeneBudget with Fixed TF–IDF. Fixed TF–IDF uses the same expression-specificity score and gene ranking as AdaGeneBudget but replaces cell-specific score-mass allocation with a shared fixed budget (Fig. S1a).

On Geneformer–Kang, AdaGeneBudget improved accuracy from 0.8786 to 0.8857, macro-F1 from 0.7130 to 0.7709, and balanced accuracy from 0.6827 to 0.7563. The two methods used comparable realized query budgets, with 404.2 genes per cell for AdaGeneBudget and 390.3 for Fixed TF–IDF. Because the two methods share the same expression-specificity ranking, this comparison is consistent with an additional contribution from adaptive budget allocation beyond the ranking rule itself.

The adaptive policy also produced distinct budget distributions across cell types (Fig. S1b). Megakaryocytes and several lymphoid populations frequently received budgets below the matched fixed value, whereas monocyte and dendritic populations more often received larger budgets. This behavior follows from the cumulative score-mass criterion: cells with more concentrated score distributions reach the target mass using fewer genes, whereas cells with more diffuse distributions require larger budgets.

**Hyperparameter sensitivity.** We evaluated sensitivity to the three budget parameters $\tau$, $K_{\min}$, and $K_{\max}$ across all four backbone–dataset pairs (Fig. S1c–f), using the same default configuration $(\tau, K_{\min}, K_{\max}) = (0.90, 128, 600)$ without pair-specific tuning.

On scGPT–Kang, macro-F1 ranged from 0.823 to 0.870 across the tested configurations, with more restrictive settings such as $K_{\max} = 300$ generally reducing performance. The default configuration achieved a macro-F1 of 0.854 and remained competitive with neighboring settings.

For the remaining three backbone–dataset pairs, the frozen default achieved macro-F1 values of 0.8932, 0.7709, and 0.7727 on scGPT–PBMC, Geneformer–Kang, and Geneformer–PBMC, respectively. These values were within 0.19, 0.69, and 0.56 percentage points of the best configurations in the corresponding tested grids. Sensitivity was setting dependent: scGPT–PBMC and Geneformer–PBMC were comparatively stable across the evaluated grid, whereas Geneformer–Kang showed larger degradation under some restrictive budget combinations. Overall, the default configuration remained close to the best tested setting across all evaluated backbone–dataset pairs without pair-specific tuning.

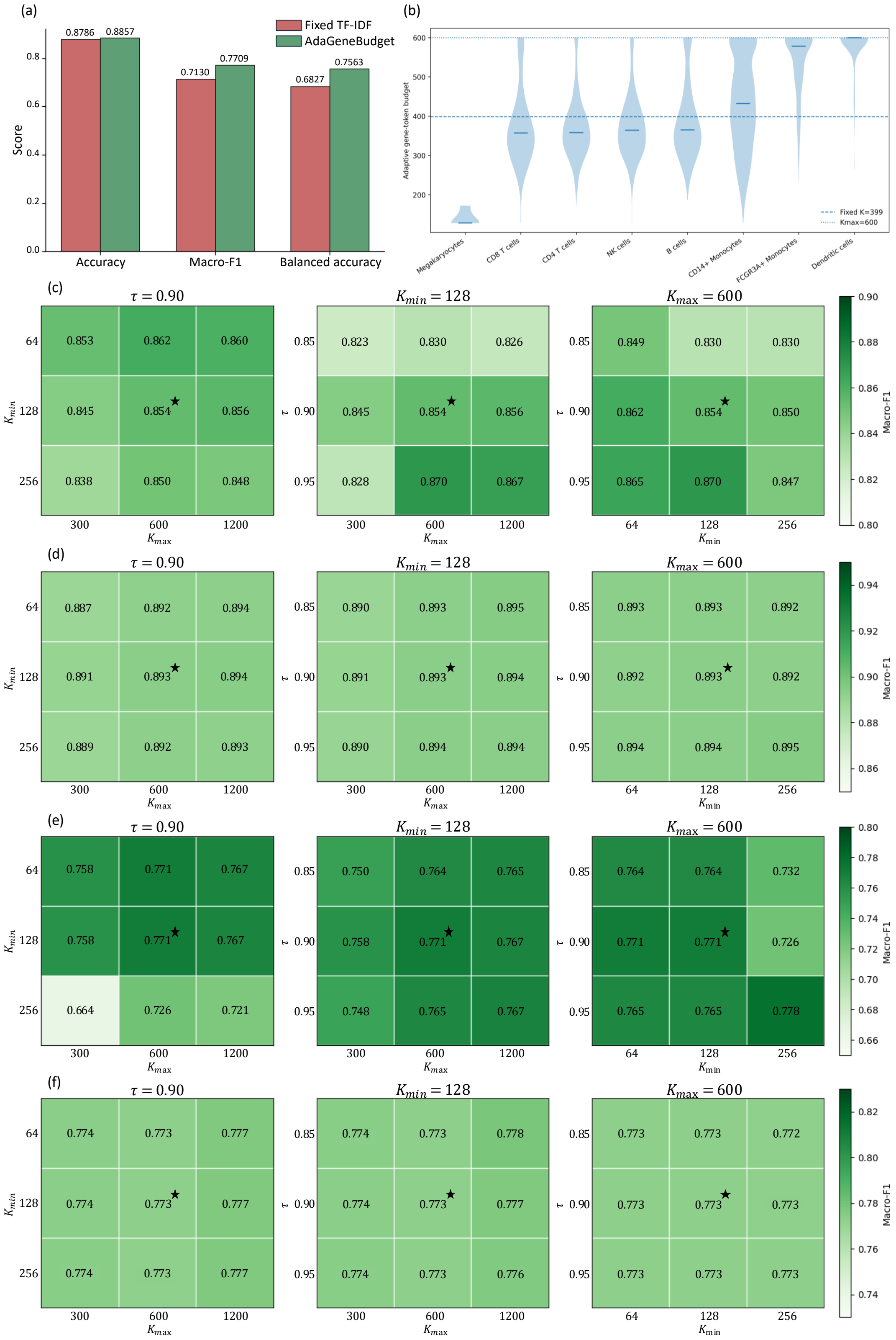


Fig. S1. **Ablation and hyperparameter sensitivity.** (a) Annotation performance of Fixed TF–IDF and AdaGeneBudget on Geneformer–Kang under comparable realized gene budgets. (b) Cell-type-specific distributions of the adaptive gene-token budget on scGPT–Kang; horizontal lines indicate the matched fixed budget ($K = 399$) and maximum adaptive budget ($K_{\max} = 600$). (c–f) Macro-F1 sensitivity to $\tau$, $K_{\min}$, and $K_{\max}$ on (c) scGPT–Kang, (d) scGPT–PBMC, (e) Geneformer–Kang, and (f) Geneformer–PBMC, respectively. The default configuration is $(\tau, K_{\min}, K_{\max}) = (0.90, 128, 600)$. Stars indicate the default configuration.

#### *S2. Statistical Uncertainty*

We quantified sampling uncertainty using 2,000 paired, class-stratified bootstrap resamples of held-out query cells. Within each bootstrap replicate, query cells were resampled with replacement within cell type, and the same resampled indices were applied to all methods. For stochastic selection policies, metrics were computed from the five saved selection-seed predictions and averaged within each replicate; deterministic methods used their single prediction artifact. We report 95% percentile confidence intervals. These intervals quantify query-cell sampling uncertainty within the held-out donor.

**Annotation performance.** Table S1 reports bootstrap confidence intervals for AdaGeneBudget across all four backbone–dataset pairs. Table S2 reports paired macro-F1 differences between AdaGeneBudget and the corresponding comparators.

Table S1. Bootstrap uncertainty for AdaGeneBudget annotation performance. Intervals are 95% percentile confidence intervals from 2,000 paired, class-stratified bootstrap resamples of held-out query cells.

| Backbone | Dataset | Accuracy [95% CI] | Macro-F1 [95% CI] | Balanced acc. [95% CI] |
|---|---|---|---|---|
| scGPT | Kang | 0.9493 [0.9430, 0.9552] | 0.8538 [0.8097, 0.8972] | 0.8685 [0.8044, 0.9109] |
| scGPT | PBMC | 0.9338 [0.9308, 0.9368] | 0.8932 [0.8879, 0.8985] | 0.8771 [0.8713, 0.8827] |
| Geneformer | Kang | 0.8857 [0.8774, 0.8939] | 0.7709 [0.7169, 0.8079] | 0.7563 [0.6892, 0.8008] |
| Geneformer | PBMC | 0.8434 [0.8392, 0.8477] | 0.7727 [0.7668, 0.7787] | 0.7758 [0.7703, 0.7815] |

Table S2. Paired bootstrap differences in macro-F1 between AdaGeneBudget and comparison policies. Positive values favor AdaGeneBudget. Intervals are 95% percentile confidence intervals from 2,000 paired bootstrap resamples.

| Backbone | Dataset | Comparator | AdaGeneBudget − comparator | 95% CI |
|---|---|---|---|---|
| scGPT | Kang | Native Bucketed | +0.0132 | [−0.0204, +0.0673] |
| | | Matched Random | +0.0224 | [−0.0056, +0.0608] |
| | | Top-Expression | +0.0199 | [−0.0207, +0.0739] |
| scGPT | PBMC | Native Bucketed | +0.0191 | [+0.0150, +0.0232] |
| | | Matched Random | +0.0587 | [+0.0542, +0.0628] |
| | | Top-Expression | +0.0071 | [+0.0031, +0.0113] |
| Geneformer | Kang | Native Bucketed | +0.1095 | [+0.0561, +0.1471] |
| | | Native Rank | +0.0697 | [+0.0086, +0.1424] |
| | | Matched Random | +0.1804 | [+0.1275, +0.2172] |
| | | Top-Expression | +0.0618 | [+0.0019, +0.1346] |
| Geneformer | PBMC | Native Bucketed | +0.0042 | [−0.0023, +0.0106] |
| | | Native Rank | +0.0170 | [+0.0102, +0.0242] |
| | | Matched Random | +0.1529 | [+0.1468, +0.1598] |
| | | Top-Expression | +0.0130 | [+0.0067, +0.0197] |

Overall, the paired bootstrap analysis supported the larger macro-F1 advantages of

AdaGeneBudget, particularly against compressed baselines. Confidence intervals for several smaller differences included zero, indicating that these numerical advantages should be interpreted descriptively.

**Biological-preservation metrics.** For Geneformer–Kang stimulation-associated pathway retention, expressing query cells were resampled with replacement separately for each pathway gene before recomputing gene-level conditional retention and pathway-level means. For Geneformer–PBMC lineage-marker coverage, query cells were resampled with replacement within lineage before recomputing lineage-level coverage. The same resampled cells were applied across methods in each replicate, and Matched Random was averaged across its five saved selection seeds.

Table S3. Bootstrap uncertainty for biological-preservation metrics. Intervals are 95% percentile confidence intervals from 2,000 paired bootstrap resamples of held-out query cells.

| Dataset / metric | Method | Point estimate | 95% CI |
|---|---|---|---|
| Geneformer–Kang pathway retention | Native Bucketed | 1.0000 | [1.0000, 1.0000] |
| | Native Rank | 0.6723 | [0.6660, 0.6785] |
| | Matched Random | 0.6143 | [0.6106, 0.6181] |
| | AdaGeneBudget | 0.8066 | [0.8044, 0.8089] |
| Geneformer–PBMC lineage-marker coverage | Native Bucketed | 0.9160 | [0.9143, 0.9177] |
| | Native Rank | 0.5894 | [0.5870, 0.5920] |
| | Matched Random | 0.2635 | [0.2621, 0.2647] |
| | AdaGeneBudget | 0.8196 | [0.8178, 0.8215] |

For the biological-preservation metrics, AdaGeneBudget retained a clear bootstrap advantage over the compressed comparison policies, while Native Bucketed remained the uncompressed reference.

#### *S3. Representation–Geometry Fidelity*

To directly assess representation fidelity under gene-token compression, we compared held-out query embeddings with same-backbone reference embeddings. For scGPT, deterministic Full Bucketed embeddings were used as the reference; for Geneformer, Native Bucketed embeddings were used. We evaluated same-cell cosine similarity and exact nearest-neighbor Recall@10 to measure embedding displacement and local-neighborhood preservation.

To test whether neighborhood preservation was driven only by broad cell-type separation, we additionally recomputed neighborhood overlap and pairwise similarity ordering within each cell type. Within each cell type, Recall@10 was computed as the mean overlap between each cell's exact 10-nearest-neighbor sets in the reference and compressed embeddings. Within-type $\rho$ was computed as the Spearman correlation between pairwise cosine similarities in the two embedding spaces, using up to 1,000 query cells per cell type for computational tractability. These metrics were macro-averaged over cell types with at least 11 query cells so that exact $k = 10$ neighborhoods were well defined. These within-cell-type results were macro-averaged over cell types with at least 11 query cells so that exact $k = 10$ neighborhoods were well defined.

Table S4. Representation fidelity relative to same-backbone reference embeddings. Within-cell-type metrics are macro-averaged over cell types with at least 11 query cells. Higher values indicate greater preservation.

| Backbone | Dataset | Method | Same-cell cosine | Global R@10 | Within-type R@10 | Within-type $\rho$ |
|---|---|---|---|---|---|---|
| scGPT | Kang | Native | 0.9957 | 0.3428 | 0.4575 | 0.9126 |
| | | Random | 0.9917 | 0.2053 | 0.3123 | 0.8407 |
| | | AdaGeneBudget | 0.9746 | 0.2310 | 0.3439 | 0.8557 |
| | PBMC | Native | 0.9971 | 0.2032 | 0.3152 | 0.9423 |
| | | Random | 0.9921 | 0.0842 | 0.1595 | 0.8460 |
| | | AdaGeneBudget | 0.8971 | 0.1461 | 0.2057 | 0.8799 |
| Geneformer | Kang | Native Rank | 0.9603 | 0.2747 | 0.3664 | 0.7027 |
| | | Random | 0.9837 | 0.2309 | 0.3191 | 0.7947 |
| | | AdaGeneBudget | 0.9781 | 0.2914 | 0.4279 | 0.9002 |
| | PBMC | Native Rank | 0.8925 | 0.1145 | 0.2132 | 0.8349 |
| | | Random | 0.9265 | 0.0247 | 0.0869 | 0.5389 |
| | | AdaGeneBudget | 0.8463 | 0.0729 | 0.1466 | 0.7936 |

AdaGeneBudget achieved higher global and within-cell-type neighborhood Recall@10 than token-matched Random in all four backbone–dataset pairs. Within-cell-type pairwise similarity correlations showed the same consistent ordering. Thus, the local-structure advantage of expression-specificity-based selection was not explained solely by preservation of broad cell-type separation. Same-cell cosine similarity did not always follow the same ordering, indicating that small embedding displacement alone does not guarantee preservation of local cellular neighborhoods.

We further used cellular structure derived from the complete expression matrix as a complementary reference, rather than assuming that the full/native foundation-model embedding itself fully represents transcriptomic structure. For each dataset, the held-out query-cell expression matrix was library-size normalized to 10,000 counts per cell, log-transformed with $\log(1 + x)$, and projected to 50 dimensions using TruncatedSVD without prior feature selection. The resulting expression-space representation was shared across backbones for the same dataset.

We quantified agreement with this expression-space reference using two complementary metrics. Expression R@10 denotes the mean overlap between each cell's exact 10-nearest neighbors in expression space and in the corresponding foundation-model embedding. Expression $\rho$ denotes the Spearman correlation between pairwise cosine similarities in expression space and those in the model embedding, computed on a fixed subsample of up to 2,000 held-out query cells. These metrics therefore assess local-neighborhood and broader pairwise-geometry agreement with the complete-expression representation, respectively.

AdaGeneBudget also exceeded token-matched Random in both expression-referenced metrics for all four backbone–dataset pairs. Notably, even the full/native foundation-model references showed limited exact neighborhood overlap with the expression-space reference, indicating that foundation-model and transcriptomic geometries are related but not identical. We

Table S5. Agreement with cellular structure derived from the complete expression matrix. "Reference" denotes Full Bucketed for scGPT and Native Bucketed for Geneformer. Higher values indicate greater agreement with the expression-space reference.

| Backbone | Dataset | Method | Expression R@10 | Expression $\rho$ |
|---|---|---|---|---|
| scGPT | Kang | Reference | 0.1264 | 0.8471 |
| | | Random | 0.0848 | 0.8082 |
| | | AdaGeneBudget | 0.1209 | 0.8427 |
| | PBMC | Reference | 0.1523 | 0.8179 |
| | | Random | 0.0455 | 0.8058 |
| | | AdaGeneBudget | 0.1123 | 0.8720 |
| Geneformer | Kang | Reference | 0.0878 | 0.7682 |
| | | Random | 0.0507 | 0.7501 |
| | | AdaGeneBudget | 0.0817 | 0.7924 |
| | PBMC | Reference | 0.0450 | 0.7434 |
| | | Random | 0.0102 | 0.5973 |
| | | AdaGeneBudget | 0.0370 | 0.7402 |

therefore interpret these results as complementary evidence of structural preservation rather than treating either representation as a ground-truth geometry.

### *S4. Reference-Dependence Analysis*

We evaluated the sensitivity of AdaGeneBudget to the reference cohort used to estimate the inverse detection-frequency (IDF) weights on Geneformer–Kang. Only the IDF-fitting reference was varied; the held-out Kang query set, full Kang reference embedding bank, Geneformer checkpoint, and AdaGeneBudget hyperparameters were kept fixed.

**Reference-size reduction.** Starting from the canonical 19,583-cell Kang reference cohort,[1] we constructed 50% and 25% IDF-fitting subsets while retaining all seven reference donors and both conditions through donor-by-condition stratified subsampling. Five independently sampled subsets were evaluated for each fraction. The resulting IDF vectors remained highly correlated with the canonical weights, while downstream annotation performance and the realized query-cell budgets remained close to those obtained using the full reference (Table S6).

**Reference-donor composition.** To evaluate sensitivity to donor composition, we recomputed the IDF weights after excluding each of the seven reference donors in turn. Across the seven leave-one-reference-donor-out settings, macro-F1 was $0.7694 \pm 0.0032$, compared with 0.7709 using the full canonical reference. Detailed donor-level results are shown in Table S7.

**Cross-cohort reference mismatch.** We further evaluated a deliberately mismatched reference by estimating IDF weights from the 132,137-cell PBMC reference cohort[2] while retaining the Kang query set and the full Kang reference embedding bank for downstream mapping. Among the 12,288 Geneformer-supported Kang genes, 12,149 (98.87%) were directly represented in the PBMC reference. For the remaining 139 genes, we assigned the median PBMC IDF value rather than treating them as completely undetected and assigning an artificially large rarity weight. The PBMC-derived IDF vector had a Pearson correlation of 0.8291 with the canonical Kang IDF vector. Despite this change in the weighting statistics, macro-F1 did not decrease in this experiment (0.7869 versus 0.7709 with the canonical Kang reference).

Table S6. Reference-dependence analysis on Geneformer–Kang. Only the cohort used to estimate the IDF weights was varied; the held-out query set, full Kang reference embedding bank, Geneformer checkpoint, and AdaGeneBudget hyperparameters were kept fixed. For the 50% and 25% settings, downstream performance metrics and realized query-cell budgets are reported as mean ± standard deviation across five independently sampled donor-by-condition stratified subsets; IDF correlation is reported as the mean across the same subsets. The leave-one-donor-out row summarizes the seven donor exclusions.

| IDF-fitting reference | Reference cells | IDF corr. vs canonical | Accuracy | Macro-F1 | Balanced acc. | Test mean genes |
|---|---|---|---|---|---|---|
| Canonical Kang (100%) | 19,583 | 1.0000 | 0.8857 | 0.7709 | 0.7563 | 404.2 |
| Kang 50% | 9,791 | 0.9966 | 0.8857 ± 0.0023 | 0.7688 ± 0.0066 | 0.7540 ± 0.0083 | 404.2 ± 0.04 |
| Kang 25% | 4,895 | 0.9919 | 0.8871 ± 0.0020 | 0.7696 ± 0.0056 | 0.7564 ± 0.0063 | 404.2 ± 0.06 |
| Leave-one-reference-donor-out | 15,003–18,541 | 0.9990 | 0.8870 ± 0.0025 | 0.7694 ± 0.0032 | 0.7563 ± 0.0032 | ≈ 404.2 |
| PBMC mismatched reference | 132,137 | 0.8291 | 0.8971 | 0.7869 | 0.7838 | 383.1 |

Table S7. Leave-one-reference-donor-out analysis on Geneformer–Kang. Each row excludes one Kang reference donor when estimating the IDF weights; all other components of the evaluation are unchanged.

| Excluded donor | Remaining cells | IDF corr. vs canonical | Accuracy | Macro-F1 | Balanced acc. | Test mean genes |
|---|---|---|---|---|---|---|
| patient_101 | 17,559 | 0.99937 | 0.8904 | 0.7710 | 0.7563 | 404.2 |
| patient_1016 | 16,225 | 0.99877 | 0.8847 | 0.7641 | 0.7501 | 404.0 |
| patient_1039 | 18,481 | 0.99948 | 0.8890 | 0.7742 | 0.7608 | 404.2 |
| patient_107 | 18,541 | 0.99967 | 0.8892 | 0.7696 | 0.7561 | 404.2 |
| patient_1244 | 16,240 | 0.99886 | 0.8855 | 0.7693 | 0.7579 | 404.4 |
| patient_1256 | 15,449 | 0.99850 | 0.8837 | 0.7671 | 0.7561 | 404.2 |
| patient_1488 | 15,003 | 0.99838 | 0.8868 | 0.7704 | 0.7565 | 404.2 |

Overall, these results suggest limited sensitivity, within the tested conditions, to moderate reductions in reference size, changes in donor composition, and cross-cohort weight estimation.

#### *S5. External Validation on Triple-Negative Breast Cancer*

To assess whether the efficiency–utility tradeoff observed in the primary blood-focused experiments extends to distinct tissues and disease settings, we performed additional donor-disjoint validation on a triple-negative breast-cancer (TNBC) dataset.

One donor was held out as the query cohort and all remaining donors formed the reference cohort. Expression-specificity weights and matched fixed budgets were estimated using reference cells only, without using query statistics. We retained the default AdaGeneBudget hyperparameters $(\tau, K_{\min}, K_{\max}) = (0.90, 128, 600)$ without dataset-specific tuning and evaluated the same frozen scGPT whole-human and Geneformer V1-10M checkpoints used in the primary experiments. Query cells were annotated by exact cosine 5-nearest-neighbor majority voting against the corresponding reference embedding bank.

Table S8. Computational efficiency on the TNBC cohort. Speedup is measured relative to Native Bucketed within each backbone.

| Backbone | Dataset | Input policy | Mean genes ↓ | Throughput (cells/s) ↑ | Speedup | Peak memory (GB) ↓ |
|---|---|---|---|---|---|---|
| scGPT | TNBC | Native Bucketed | 1100.8 | $481.7 \pm 2.3$ | 1.00× | 1.376 |
| | | Matched Random | 576.0 | $1012.5 \pm 6.8$ | 2.10× | 0.775 |
| | | Top-Expression | 576.0 | $896.2 \pm 2.3$ | 1.86× | 0.775 |
| | | AdaGeneBudget | 581.2 | $867.6 \pm 6.9$ | 1.80× | 0.791 |
| Geneformer | TNBC | Native Bucketed | 1343.5 | $189.3 \pm 1.3$ | 1.00× | 13.527 |
| | | Native Rank | 584.2 | $712.6 \pm 2.4$ | 3.77× | 3.925 |
| | | Matched Random | 584.2 | $729.9 \pm 3.5$ | 3.86× | 3.925 |
| | | Top-Expression | 584.2 | $722.0 \pm 4.2$ | 3.81× | 3.925 |
| | | AdaGeneBudget | 585.6 | $654.7 \pm 7.3$ | 3.46× | 3.977 |

**Dataset and donor-disjoint split.** The treatment-naive TNBC dataset[3] contains 427,823 cells from 101 donors and 32,354 genes (CELLxGENE dataset `6f9de485-58cd-4342-bfc4-b3d3dd223aa8`). We held out donor P74 as the query cohort, leaving 418,671 cells from the remaining donors as the reference cohort and 9,152 P74 cells as queries. P74 was absent from the reference cohort.

P74 was selected as the largest audited donor containing all seven coarse cell types, with all query coarse types represented in the remaining donors and with adequate per-class support. The dataset contains seven coarse `cell_type` classes and 61 dataset-wide `cell_state` labels. The P74 query cohort contains 54 cell states, all represented in the reference cohort, and 35 of these states have at least 20 query cells.

For scGPT, genes were matched using `feature_name`, yielding 23,713 matched genes. For Geneformer, Ensembl identifiers were matched using the Genecorpus-30M dictionaries, yielding 19,907 supported genes. IDF weights were estimated from the 418,671 non-P74 reference cells only. The reference-fitted matched fixed budgets were $K = 583$ for scGPT and $K = 592$ for Geneformer.

**Fine-grained annotation protocol.** We first evaluated the dataset-provided coarse `cell_type` labels. Because this seven-class task was nearly saturated under the native representations, with Native Bucketed macro-F1 of approximately 0.996 for scGPT and 0.985 for Geneformer, we used the finer dataset-provided `cell_state` labels as the primary TNBC benchmark. The 54 query cell states provide a more discriminative evaluation of within-lineage structure than the seven broad cellular compartments.

For the primary evaluation, all 418,671 non-P74 cells formed the reference embedding bank and the 9,152 P74 cells formed the query set. Annotation used exact cosine 5-nearest-neighbor majority voting.

**Computational efficiency.** Throughput was measured over the full 427,823-cell cohort using the same timing protocol as in the primary experiments. Table S8 shows that AdaGeneBudget reduced the mean query input size from 1,100.8 to 581.2 genes for scGPT and from 1,343.5 to 585.6 genes for Geneformer. This yielded 1.80× and 3.46× throughput speedups over Native Bucketed, respectively.

**Fine-grained annotation utility.** Table S9 reports performance using the 54 `cell_state` labels represented in the held-out P74 query cohort.

For scGPT, AdaGeneBudget achieved macro-F1 of 0.5411, compared with 0.5396 for Native Bucketed, 0.5283 for Top-Expression, and 0.4885 for Matched Random. For Geneformer, AdaGeneBudget achieved macro-F1 of 0.3392, compared with 0.3226 for Native Bucketed, 0.3066 for Top-Expression, 0.2954 for Native Rank, and 0.1972 for Matched Random. AdaGeneBudget achieved the highest numerical accuracy, macro-F1, and balanced accuracy among the evaluated policies for both backbones in this fine-grained TNBC evaluation.

Table S9. Donor-held-out fine-grained cell-state annotation performance on the TNBC cohort. Stochastic policies are reported as mean ± standard deviation across selection seeds; deterministic policies are reported once.

| Backbone | Dataset | Input policy | Mean genes | Accuracy ↑ | Macro-F1 ↑ | Balanced accuracy ↑ |
|---|---|---|---|---|---|---|
| scGPT | TNBC | Native Bucketed | 1100.8 | $0.6584 \pm 0.0026$ | $0.5396 \pm 0.0053$ | $0.5691 \pm 0.0054$ |
| | | Matched Random | 576.0 | $0.5961 \pm 0.0041$ | $0.4885 \pm 0.0047$ | $0.5217 \pm 0.0058$ |
| | | Top-Expression | 576.0 | 0.6488 | 0.5283 | 0.5578 |
| | | AdaGeneBudget | 581.2 | 0.6639 | 0.5411 | 0.5743 |
| Geneformer | TNBC | Native Bucketed | 1343.5 | 0.4079 | 0.3226 | 0.3598 |
| | | Native Rank | 584.2 | 0.3821 | 0.2954 | 0.3501 |
| | | Matched Random | 584.2 | $0.3062 \pm 0.0025$ | $0.1972 \pm 0.0039$ | $0.2302 \pm 0.0097$ |
| | | Top-Expression | 584.2 | 0.4083 | 0.3066 | 0.3421 |
| | | AdaGeneBudget | 585.6 | 0.4300 | 0.3392 | 0.3729 |

Overall, AdaGeneBudget consistently outperformed token-matched Random in annotation utility while retaining substantial computational gains relative to Native Bucketed. This experiment extends the evaluation beyond the original blood-focused settings to breast-tumor data. On TNBC, AdaGeneBudget achieved the highest numerical annotation performance among the evaluated policies for both scGPT and Geneformer. These results provide evidence that the proposed gene-token selection strategy remains effective in an additional tissue and disease setting, while also showing that the relative advantage of adaptive selection is not uniform across datasets and backbones.

### *S6. Noise–Sensitivity Analysis*

Because inverse detection-frequency weighting assigns larger weights to genes detected in fewer reference cells, we examined whether low-frequency features could receive disproportionate priority. We performed this analysis on scGPT–PBMC. Low-frequency genes were defined as those detected in at most 1% of the reference cohort. For each eligible expressed gene–cell pair in the held-out query set, we recorded whether the gene was retained by AdaGeneBudget. We additionally compared AdaGeneBudget with an expression-only control that selected the highest-expressed genes using the same realized cell-specific budget $K_i$, thereby isolating the contribution of inverse-frequency weighting from the number of retained genes. Within each query cell, weakly expressed genes were defined as those in the bottom quartile of positive expression values.

AdaGeneBudget therefore preferentially retained low-frequency genes beyond what could

Table S10. Noise-sensitivity analysis on scGPT–PBMC. Low-frequency genes were defined by reference detection frequency $\leq 1\%$, and weak expression by the bottom quartile of positive expression within each query cell.

| Quantity | Value |
|---|---|
| Low-frequency pairs among eligible gene–cell pairs | 0.695% |
| Low-frequency pairs among Ada-selected pairs | 2.09% |
| Enrichment among selected inputs | $\approx 3.0\times$ |
| Selection rate among low-frequency, weak-expression pairs: Ada / expression-only | 68.5% / 2.1% |
| Selected TF–IDF score mass from low-frequency genes | 1.12% |

be explained by expression magnitude alone, including weakly expressed low-frequency features. However, low-frequency features remained a small absolute component of the selected input, comprising 2.09% of selected gene–cell pairs and 1.12% of the selected TF–IDF score mass. Because low reference frequency does not itself establish that a feature is a technical artifact, this analysis does not directly measure artifact retention. Rather, it indicates a potential sensitivity to rare technical artifacts, without suggesting that low-frequency features dominate the selected representation in this setting.
